# Big dairy data to disentangle the effect of geo-environmental, physiological and morphological factors on milk production of mountain-pastured Braunvieh cows

**DOI:** 10.1101/2020.04.16.042028

**Authors:** Solange Duruz, Elia Vajana, Alexander Burren, Christine Flury, Stéphane Joost

**Author notes:** Laboratory of Geographic Information System (LASIG), Ecole polytechnique Fédérale de Lausanne (EPFL), 1015 Lausanne, Switzerland. This author should be considered as co-senior author.

## Abstract

The transhumance system, which consists in moving animals to high mountain pastures during summer, plays a considerable role in preserving both local biodiversity and traditions, as well as protecting against natural hazard. In cows, particularly, milk production is observed to decline as a response to food shortage and climatic stress, leading to atypical lactation curves that are barely described by current lactation models. Here, we relied on five million monthly milk records from over 200,000 Braunvieh and Original Braunvieh cows to devise a new model accounting for transhumance, and test the influence of environmental, physiological, and morphological factors on cattle productivity. Counter to expectations, environmental conditions in the mountain showed a globally limited impact on milk production during transhumance, with cows in favourable conditions producing only 10% less compared to cows living in adverse conditions, and with precipitation in spring and altitude revealing to be the most production-affecting variables. Conversely, physiological factors as lactation number and pregnancy stage presented an important impact over the whole lactation cycle with 20% difference in milk production, and may therefore alter the way animals respond to transhumance. Finally, the considered morphological factors (cow height and foot angle) presented a smaller impact during the whole lactation cycle (10% difference in milk production). The present findings can help farmers to establish sustainable strategies for alleviating the negative effects of transhumance on productivity and preserving this important livestock practice.

## 2. Introduction

Transhumance, which consists in moving livestock to high mountain pastures in the summer months, provides both ecological and socio-cultural services to the human populations living in the mountainous regions of many European countries[1–3]. Indeed, transhumance-annexed grazing sustains and preserves endemic plant communities [4], feed local cattle to produce traditional alpine cheese, and attract many tourism-related activities [5]. Further, it counteracts land abandonment in mountain areas and therefore contributes preserving landscape against scrubs growth and vegetation encroachment [6], as well as natural hazards such as avalanches [7] and wild fires [5]. The term “alping” (a translation of the German word “Alpung” or its French equivalent “montée à alpage”) will be used here to describe the approximately 100 days that dairy cattle spend on alpine pastures during the summer months. Similarly, animals brought to mountain pastures will be referred to as “alped” cows, and the alpine summer pastures will be called “alps”.

Despite such ecological and social benefits, the surface dedicated to alping decreases each year (~2400 ha per year [8]), and a questionnaire-based study revealed in 2010 that one third of the participating breeders intend to probably abandon the transhumance practice in the following decades. In summer 2018, 107’000 dairy cows were alped in Switzerland during approximately 100 days [9]. A steep drop in milk production is observed during this period, which hampered the evaluation of lactation curves through standard models that assume a linear decrease in production [10] after the maximum milk yield is reached (i.e. ~100 days after calving) [11]. Among the explanations proposed to interpret such a detrimental effect on productivity are the food deficit intake due to the meagre grassland as found in high alpine pastures, as well as the need to tackle environmental stress due to new and sometimes harsh habitat conditions [12]. On the other hand, milk composition is known to change during alping [13,14] and results in the production of highly valuable milk products such as butter and alp cheese.

Milk production and quality is notoriously affected by a wide variety of environmental factors, including calving season, vegetation types composing animals’ diet [15–18]. Environmental temperature is also known to directly affect cattle productivity because of heat [17] or cold [19] stress. Furthermore milk quality and production of alped cows are expected to be indirectly affected by global warming, as forage quality and biomass productivity of alpine sites are likely to decrease with increasing temperature and decreasing precipitation [20,21].

Despite the existence of huge databases storing monthly milk records for several European cattle breeds, no effort has been produced so far (at least to our knowledge) to exploit such an information and understand the ways alping affects milk productivity [22]. Indeed, most of the existing literature focuses on small experiments (with sample size <100) mainly restricted to compare two groups of animals in different environmental conditions, so as to investigate the potential effects of altitude [23], vegetation type [23,24], supplemental feeding [25,26], calving season [27] or breed [12,27,28]. Furthermore, no adaptation of general model of lactation curves [11] have been proposed to account for alping, which hinders a straightforward comparison of lactation curves for alped cow. Last but not least, the overall impact of environmental factors and global warming on milk production during alping is also still unknown.

For these reasons, a better understanding and characterisation of the impacts of transhumance on milk production and the way production is influenced by environmental factors is needed. To fill this gap, we relied on over five million monthly test-day milk records collected between 2000 and 2015 from more than 200,000 Braunvieh cows, a local Swiss cattle breed well adapted to the alpine pastures. Then, we used this information to: 1) devise a new mathematical model to fit lactation during alping; and 2) investigate the influence of environment on milk production, together with the effect of physiological and morphological factors on milk production during alping. This can be achieved thanks to biogeoinformatics which takes advantage of geo-referenced animal data in order to link biological and environmental information with the help of advanced informatics tools [29].

## 3. Data

### Milk records and animal information

Milk records from all alped Braunvieh cows were provided for the period 2000-2015 by the Braunvieh Schweiz AG breeding association. Importantly, a direct comparison with non-alped cows was impossible due to data unavailability, but milk measurements of alped cows entailing records from both the lowland farm and the alp, enabled the estimation of milk production in both situations. The full dataset is composed of 5,681,498 test day records (methods A4 and AT4 according to ICAR-Guidelines [30]), including 616,081 lactations derived from a total of 245,313 cows. In line with national and international rules, milk records are taken approximately on a monthly basis, with the first record taken between the 5^th^ and 42^nd^ day after calving. Each test day record included information on the following traits: Milk (kg), Fat (kg and %), Protein (kg and %), somatic cell count (1000 cells/ml), but our study specifically focused on milk production in terms of quantity (milk yield). Out of the total number of records, 1,481,387 were taken in the alps, whose altitude were systematically stored in the database, while their precise location were documented in 95% of the cases (Fig. 1). The first record in the alp is usually taken within the first four days after arrival, and is followed by three more records in the alp to encompass the entire alping period (typically 100 days). Moreover, to morphologically describe animals, linear type description and classification of cows are scored during the first lactation of all cows of the database. In our study we considered the body height at withers and the scores (1-9) for foot angle. In addition, insemination data for each lactation (date, sire’s name) are also available.

**Figure 1:**
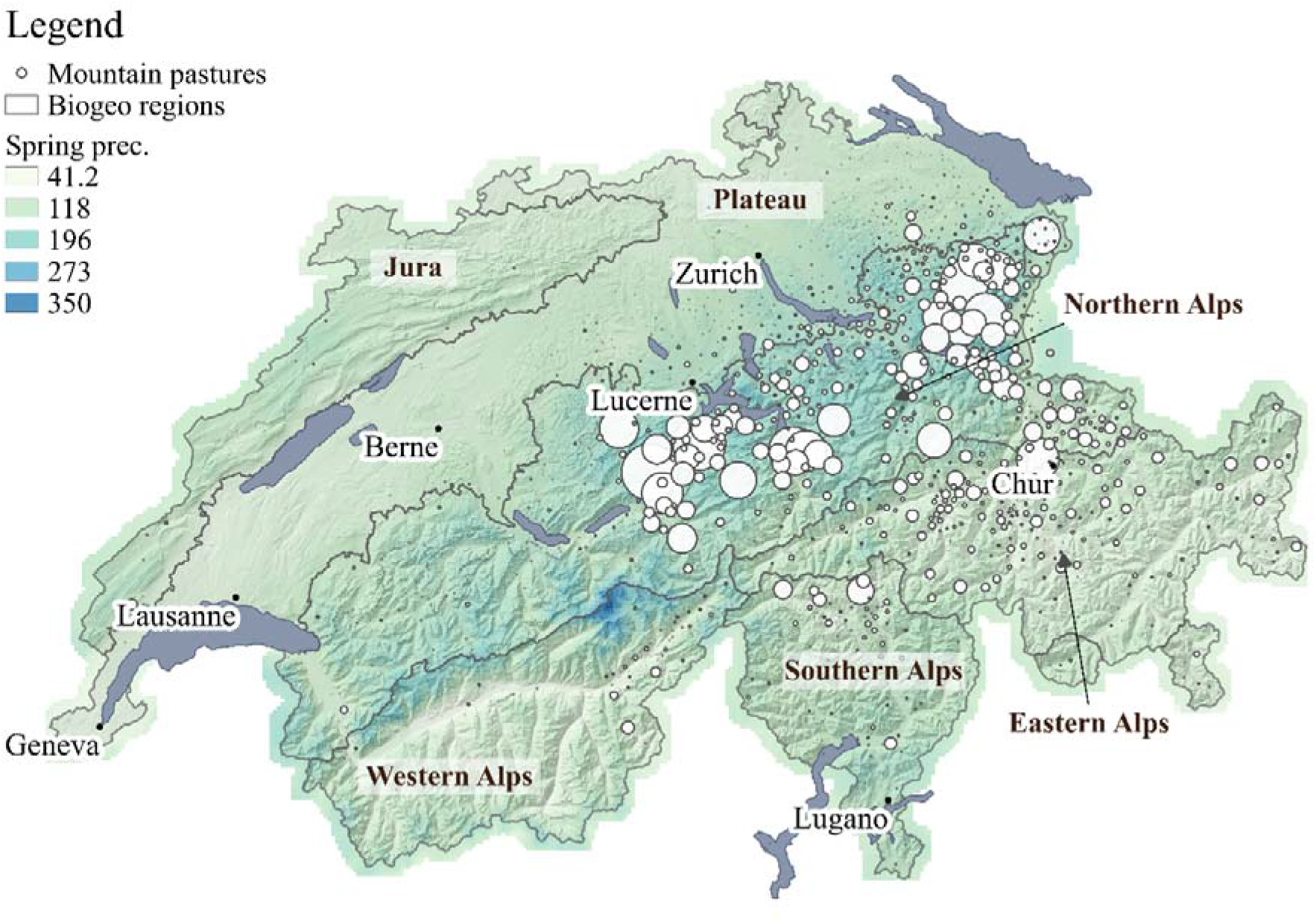
Geographic location of the alps hosting Braunvieh cows (white circles), with average monthly precipitation in mm between April and July 2015 in the background (chosen as example year). Frontiers of biogeographical regions are also reported. The size of the circles is proportional to the number of milk records taken at a given alp. The majority of the alps hosting Braunvieh cows are located in Northern and in the Eastern Alps biogeographical regions.

A stringent data quality control procedure was applied prior to analysis to remove: 1) incomplete years (which resulted in removing beginning of 2000 as well as end of 2015 due to missing lactation records); 2) cows with average interval between first and last insemination longer than 100 days (as computed over the first three lactations); 3) cows that had their first calf while being younger than two years, or older than four years; 4) cows belonging to breeds different from the Braunvieh or Original Braunvieh; 5) cows with parents other than Braunvieh or Original Braunvieh; 6) lactations shorter than 270 days; 7) lactations with calving interval shorter than 290 days; 8) lactations with alps below 1100 meters above sea level (masl) or above 2600 masl; 9) lactations with calving happening between March and August; 10) lactations from cows that had already calved more than nine times; 11) lactations with the first record taken after the 42^nd^ day after calving; 12) lactations with records taken before calving; 13) records taken before the 5^th^ day and after the 500^th^ day after calving; 14) the second alping season (i.e. final part of lactation curves) from animals that are alped twice in the same lactation. After filtering, we obtained a final dataset composed of 3,527,138 records over 371,696 lactations from 175,474 cows.

### Factors influencing milk characteristics

Milk characteristics are known to be influenced by different factors. Meaningful predictor variables were then selected according to literature review, by assuming the same factors to be relevant in both lowland and mountain conditions. As a result, climatic and environmental indices [19,31] were taken into account together with physiological (lactation number, pregnancy stage [32,33]) and morphological factors (Table 1).

**Table 1:** List of factors included in the present study with supposed influence on lactation during alping. Factor-specific cut-off values are reported in the last column. These values are used to assess factor-specific effects on lactation (see Methods for an exhaustive explanation).

|  | Name | Description | Group cut-off |
| --- | --- | --- | --- |
| Environmental | Temperature Humidity Index (3 days) | Climatic index based on temperature and humidity, averaged over 3 days before milk record at the alp. See section on climatic data | 59.4-65.4 / >65.4 |
|  | Temperature Humidity Index (30 days) | Climatic index based on temperature and humidity, averaged over 30 days before milk record at the alp. See section on climatic data. | 59.6-63.1 / >63.1 |
|  | Cold Stress Index (3 days) | Climatic index based on temperature, wind speed and precipitation, averaged over 3 days before milk record at the alp. See section on climatic data. | 960.1-1045.9 / >1045.9 |
|  | Cold Stress Index (30 days) | Climatic index based on temperature, wind speed and precipitation, averaged over 30 days before milk record at the alp. See section on climatic data. | 997-1042.9 / >1042.9 |
|  | Spring precipitation | Average monthly precipitation [mm] between April and July; computed for each year, at the location of each alp. | <120.6 / >155.3 |
|  | Biogeographical region | Only regions with sufficient sample size were retained, and therefore two categorical variables were created. See section on biogeographical regions. . | North Alp / East Alp |
|  | Altitude | Altitude [m] of the highest alp during the lactation cycle | <1600 / >1900 |
|  | Altitude difference | Difference in altitude between the highest alp and the lowland farm. | <641 / >1021 |
|  | Aspect 100m | Aspect of the alp (North/South facing) as based on 100m-resolution DEM. | 300-60 / 120-240 |
|  | Aspect 1km | Aspect of the alp (North/South facing) as based on 1km-resolution DEM. | 300-60 / 120-240 |
| Physiological | Lactation number | Number of lactations the cow experienced since birth (correlated with animal age). | 1<sup>st</sup> lact / ≥3<sup>rd</sup> lact |
|  | Pregnancy stage | Pregnancy stage [days] at the beginning of alping. | <73 days / >153 days |
| Morphological | Height of animal | Height at withers [cm] | <139 / >143 |
|  | Foot angle | A note between 1 and 9 (with 9 being the steepest). | <4 / >6 |

### Climatic data

Climate has been observed to influence milk production [34]. Consequently, maximum and mean temperature [23] as well as daily rainfall [24] were extracted from the meteoswiss Grid-Data products database. This dataset is derived by interpolation of records from several weather stations across Switzerland, and consists of 2km-resolution raster files (1km-resolution from the year 2014 and on). Further, daily average wind speed and relative humidity were obtained from respectively 440 and 495 meteoswiss weather stations. We then interpolated these values between stations to obtain a continuous representation of the variables, with a squared inverse-distance weighting (IDW) [35] within a maximum distance of 50 km.

On the basis of such environmental data, the Temperature Humidity Index (THI) and Cold Stress Index (CSI) were computed following Bryant et al. [19], in particular::

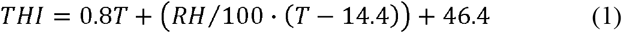

with *T* being maximum daily temperature [°C] and *RH* the relative humidity [%], and

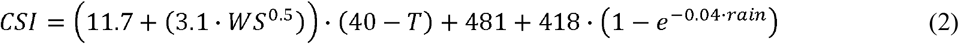

with *WS* being the wind speed [m/s], *T* the mean daily temperature [°C] and *rain* precipitation [mm]. These indices were computed over a 3- and 30-day period to account for short/long heat waves/cold spells, respectively.

### Digital Elevation Model

Due to the coarse spatial resolution of temperature data (2-km), a correction of −0.45°C/100m (i.e. the observed temperature gradient in the dataset) was applied to account for local variation in temperature due to topography. This correction was achieved using both the Digital Elevation Model DHM25 dataset produced by swisstopo [36] and the recorded altitude of the alp available in the dataset. The digital model DHM25 is a tridimensional representation of the earth’s surface in Switzerland, as based on the elevation data from the Swiss National Map 1:25,000 (NM25). A symmetric 25-m grid matrix model is then interpolated starting from the digitized contour lines and spot heights from NM25. Comparisons among control points shows an average accuracy of the produced model of 2-3 m for the pre-Alps and Alps, respectively.

### Biogeographical Region

The Federal Office for Environment (FOEN) divided Switzerland into six biogeographical regions [37], obtained using fauna and flora data and aggregating areas with common species. Species distributions being strongly related to the relief, these regions reflect in fact the topography of the country. Most of the alps hosting Braunvieh cows appear to be located in the Northern and Eastern Alps biogeographical regions. More rainfall occur in the Northern Alps when compared to the Eastern side (Fig. 1), because the mountain chain act as a barrier to precipitations coming from the West and North [38].

## 4. Methods

### Lactation curve modelling

A lactation curve is usually estimated from one single cow with repeated observations along a lactation cycle and with records taken on a daily/weekly basis [11]. Here, test-day milk records were collected monthly, making the individual-based estimates of lactation impossible because of the small number of observations with regards to the number of parameters to estimate, particularly when describing a complex curve like the one of alped cows. Therefore, we analysed several animals at a time by taking the average milk production for a given Day In Milk (DIM, or number of days after calving). Records from cows remaining at the lowland farm during the alping season (between the 15^th^ of May and the 31^st^ of August) were excluded from this average computation, while only cows at the lowland farm were considered in the average outside this time frame. Moreover, cows were grouped according to their calving month. Finally, when fitting the curve, each averaged milk yield was weighted according to the number of observations on that day.

Several models have been proposed to describe lactation curves [39], with the Wood, Wilmink, Ali-Schaeffer (AS) and Legendre polynomial formulations being the most popular [40]. Among these mathematical formulations, Wilmink proposes a linear equation that is retained in the present work given its inherent simplicity and good performance [40]. This model is written as:

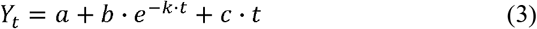

where Y_t_ is the observed variable (milk yield), t is the DIM, and a, b, c and k are the parameters to estimate. However, k is usually set to 0.1 to make this equation linear [40].

Here, we introduce additional terms to Eq. 3 in order to explicitly account for the transhumance effect. Particularly, alping has been observed to severely affect milk production, with alped animals showing a steeper linear decrease than before alping (Fig. 2). Further, alped cows usually experience a small yet rapid boost shortly after their return to the lowland farm, followed by a softer decline in milk production. Tacking these observations into account, we then propose to adapt Eq. 3 as follows:

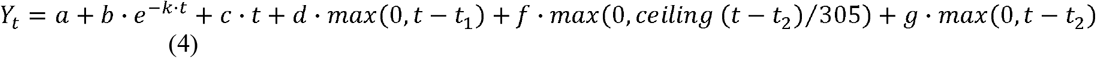

**Figure 2:**
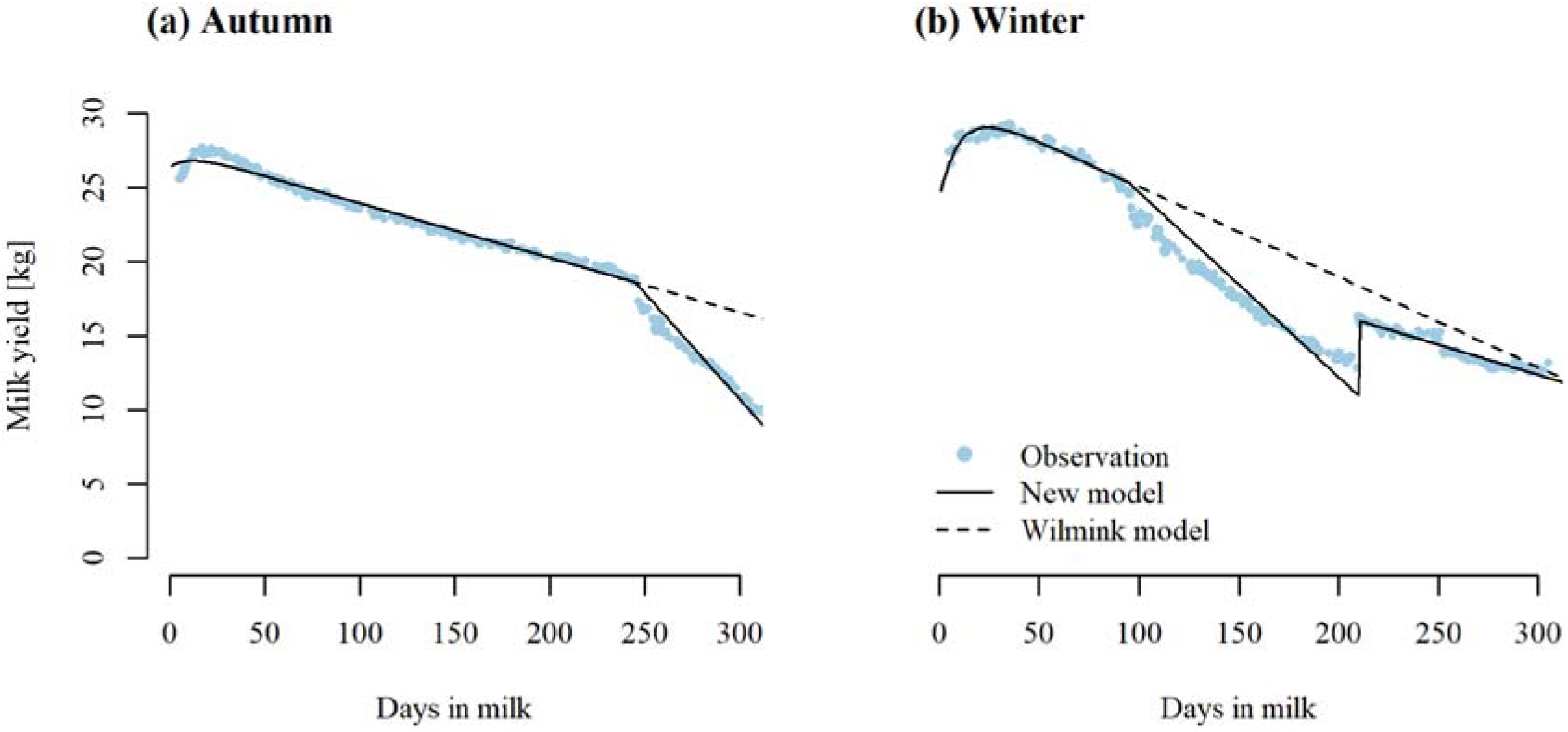
Lactation curves as derived from the proposed model (full line) and the Wilmink model (dashed line) for cows that calved in September (a) and February (b). The Wilmink model was fitted using points from the beginning of the curve only, i.e. before alping.. Each dot represents the average of milk records per day. When t>245 (a) and between 95 and 210 (b), records from the alp only are used to calculate the average, whilst records from the lowland farm only are included for the remaining time frame.

Where t_1_ is the DIM at which the cow is alped, and t_2_ is the DIM at which the cow is brought back to the lowland farm. Importantly, the expression d*max(0,t-t_1_) is the expected linear decrease during alping, so that the d-parameter reflects the effect of alping. The f*max(0,ceiling(t-t_2_)/305 captures the expected boost in production after alping and g*max(0,t-t_2_) represents the linear decrease in milk yield after alping; in the latter arguments, the max() term ensures the model to be only affected during and after alping respectively, while the ceiling expression (i.e. round to the upper integer) constructs a binary operator (0/1) to recreate the instantaneous boost after the return to the lowland farm. In our case, t_1_ and t_2_ were determined independently for each calving month. The proposed equation only works for a standard lactation period of 305 days.

The d-parameter enables the estimation of the loss in milk yield associated with alping over a given period of time. Indeed, the amount of milk lost during alping for a period of x days can be approximated with

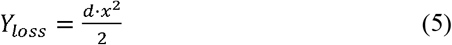

However, it is essential that the model fits well the beginning of the curve for this equation to work, which can be achieved by artificially increasing the weight of point measurements before the transhumance. Thus, weights before alping were multiplied by 100 when investigating the d-parameter depending on the calving month (Fig. 2 and 3). Furthermore, as older cows tend to calf later in the season, thereby creating a correlation between lactation number and calving month, the impact of alping according to the calving month is entangled with lactation number. Therefore, when examining milk production and the impact of alping for each calving month, only cows in their first lactation are considered (Fig. 3).

**Figure 3:**
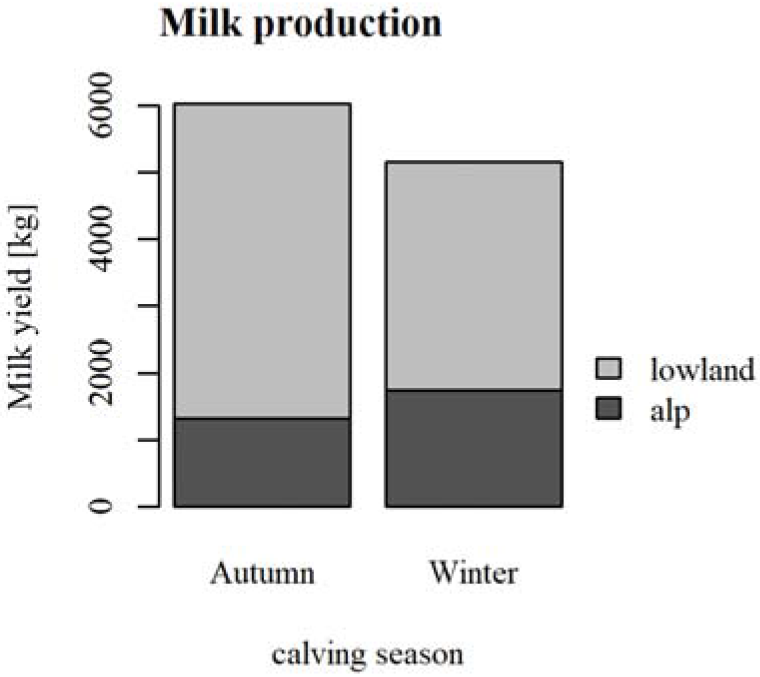
Milk production during alping (black) and from the lowland farm (grey) is reported for autumn and winter calving, as represented by the months of September and February, respectively. Only cows in their first lactation are considered here.

Ordinary linear regression models were then computed in R using the lm() function of the stats package [41] to estimate parameters in Eq. 4.

### Measuring the effect of influencing factors

For sake of interpretation, all influencing factors (i.e., explanatory variables) were grouped into environmental, physiological and morphological categories (Tab. 1). The effect of influencing factors was tested by comparing milk records produced in conditions as dissimilar as possible. Importantly, since the low number of measurements per animal imposed the use of averages, effect determination was not possible through classical regression models. Consequently, groups were created according to the first and third tertile of the distributions, in order to include animals from the most contrasted situations (environmental, physiological and morphological) while retaining enough observations to guarantee a sufficient statistical power. Since productivity is known to be optimized with mild weather conditions [34], exceptions were made for THI and CSI where the second and the third tertiles were used as the two contrast groups instead of the first and third tertile.

Group membership was assessed through the creation of a dummy variable assuming the value of 1 if belonging to the group considered, 0 otherwise. Then, the impact of influencing factors was computed by adding an interaction term to Eq. 4 that allows chosen parameters to vary as a function of the group. The here defined environmental variables affect milk production during the alping stay only. Accordingly, lactation curves were modelled only until the end of the alping season (meaning the f and g parameters not to be estimated), with the sole d-parameter varying as a function of the group. In contrast, physiological and morphological factors influence the whole lactation cycle, so that all terms of Eq. 4 (coefficients a, b, c, d, f and g) are allowed to vary as a function of the group.

Within-group production was estimated both at the lowland farm and in the alps for physiological and morphological factors or during alping season only for environmental factors, by integrating the area under the lactation curve. The between-group difference was then assessed by computing the percentage of the difference in milk production with respect to the reference group, this group being arbitrarily chosen as the one with the highest milk production during alping. The difference in the d-parameter (Δd) between the two groups is then also displayed to show how differently the concerned groups were impacted by alping. As the response differs according to the calving months, results were computed for each calving month separately and the months of September and February were chosen as representative of autumn and winter calving, respectively.

### Significance testing

Log-likelihood ratio tests were performed to investigate both the impact of adding the parameters d, f and g to the Wilmink model, and of the considered influencing factors. When testing the addition of parameters d, f and g to the Wilmink equation, Eq. 3 and 4 were considered as null and alternative models, respectively; when testing the influencing factors, the null model was constructed by removing the interaction between the dummy variable group and the parameters of Eq. 4.

The resulting G-score test-statistics were then converted into p-values, which were further corrected for multiple testing by means of the Bonferroni’s approach [42]. G-scores were evaluated using the lrtest() function from the lmtest R-package [43].

## 5. Results

### Lactation curve modelling

Overall, the proposed equation fit both the drop in milk production due to alping and the tail of the lactation curve, as illustrated here for the calving months of September and February (Fig. 2). In particular, the terms added to the Wilmink equation (Eq. 4) significantly increase the full model performance (p-value<10^-16^). In the case of autumn calving (Fig. 1a), the proposed equation fits the entire lactation cycle. For winter calving (Fig. 2b), the beginning and the end of the transhumance season appear to be the most challenging periods to be fitted because of a non-linear slope. The use of Eq. 5 can be illustrated with the autumn calving, with a d-parameter of −0.08, which is translated by a loss of 144 kg over 60 days.

Total milk production and milk production during alping is reported for the calving months of September and February (Fig. 3). For the sake of comparison among months, only cows in their first lactation are considered in this graph, as lactation number and calving month are correlated. Cows calving in autumn produce on average 6033 kg during their first lactation, among which 1320 kg are produced in the alp. In contrast, total milk production turns out to be lower for cows calving in winter (5155 kg during their first lactation), while milk production during alping is increased (1755 kg). The d-parameters for the two calving seasons being markedly different (−0.08 and −0.02 for autumn and winter calving respectively) indicates that productivity is more impacted by alping when calving occurs in autumn than when occurring in winter.

### Effect of influencing factors

The significance of the interaction between the group variable and the d-parameter is reported (Sup. Mat. S1). Hereunder, only factors with at least one calving month having a significant Δd (i.e. a significantly different impact of alping between the two contrast groups) are presented.

Among environmental conditions, THI, spring precipitation, biogeography and altitude turned out to show a significant effect on milk production during alping (Fig. 4a-l). Particularly, precipitation in spring and the biogeographical region showed the most important difference on milk production during alping, followed by altitude and altitude difference. Further, calving period appears to interact with environmental conditions, with bigger differences between groups being present in autumn.

**Figure 4:**
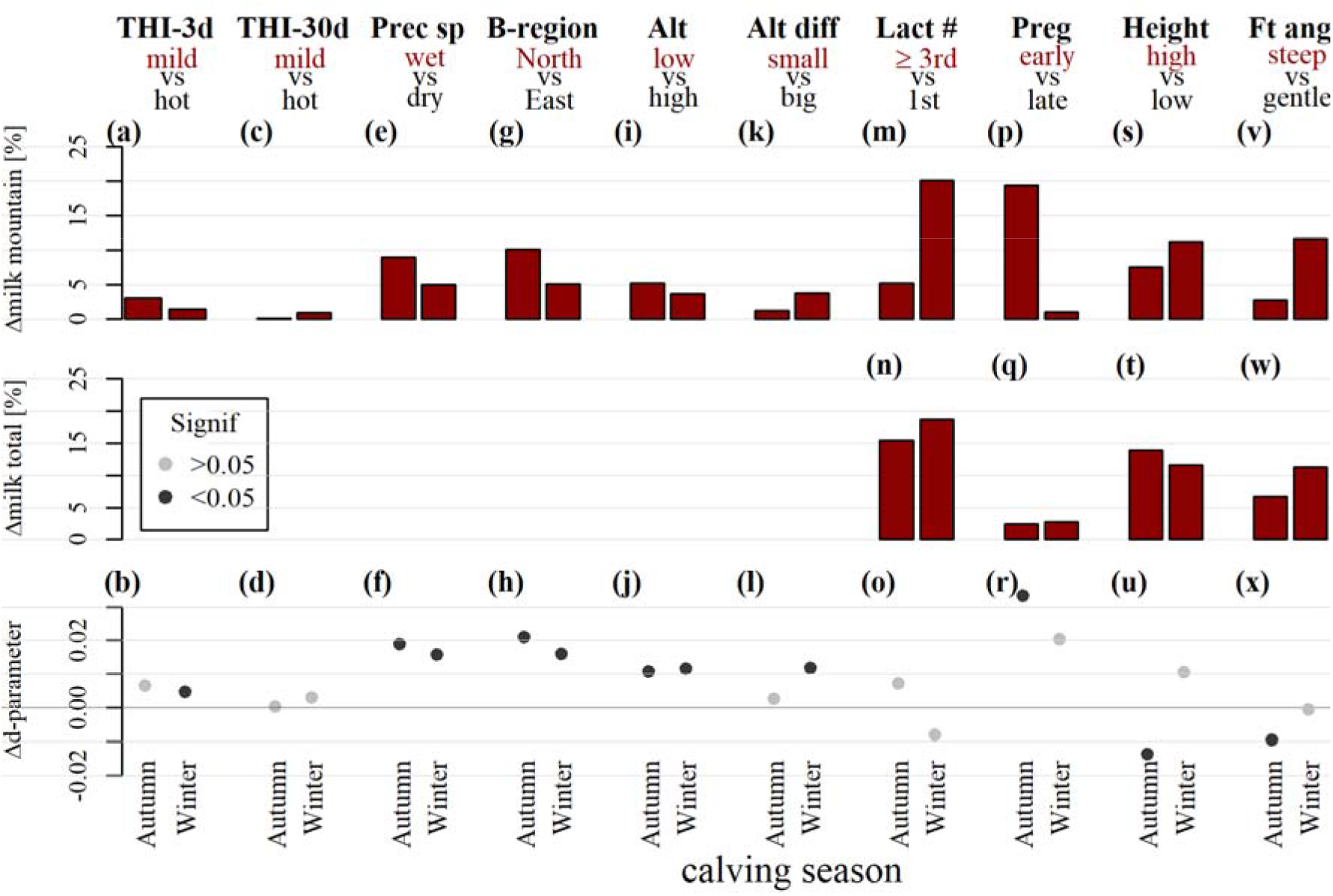
The effect of influencing factors is tested by investigating the difference in productivity between two groups of animals coming from contrasted conditions (first and third tertiles, except for THI where second and third tertile are chosen). Each factor is here reported in a separate column. At the top of each column, the factor name as well as the contrasted groups are reported; the group with highest milk yield during alping is chosen as the reference group, highlighted in red. In each barplot, the first bar shows the result for autumn calving, and the second for winter calving. The between-group difference in milk production during alping is displayed in the top panel, the between-group difference in milk production during the whole lactation in the intermediate panel, the change in the d-parameter at the bottom. The Δd-parameter indicates how the reference group is impacted by alping compared to the other group, with positive values meaning lower negative impact (see Eq. 4). Significant Δd values are plotted in black, while grey indicates non-significance. Environmental factors affects production during alping only, making a comparison of the whole milk production redundant (which is why no graph is present in the intermediate panel of the concerned variables). To facilitate the understanding of this graph, the example of lactation (Lact #) is detailed here, where we refer to cows in their third or higher lactation as third lactation cows: third lactation cows produce 5% more milk during alping than first lactation cows when calving in autumn and even 20% more when calving in winter (m). When considering the whole lactation, third lactation cows produce 15% more milk than first lactation cows when calving in autumn and 19% when calving in winter (n). Third lactation cows calving in autumn are slightly less negatively impacted by alping (positive Δd) than first lactation cows; an inverse behaviour is observed for winter calving, although both relationships are not significant (o). THI: Temperature Humidity Index, as averaged over 3 (THI-3d) or 30 (THI-30d) days. Prec sp: precipitation in spring. B-region: biogeographical region. Alt (diff): (difference in) altitude, lact #: lactation number, Preg: pregnancy stage, height: height at withers and ft ang: foot angle.

The effect of environmental factors are small compared to those of physiological factors, where the biggest effect is found for pregnancy stage for winter calving with a difference in milk production during alping of 20%. Although third and higher lactation cows produce more milk during the whole lactation cycle including alping (Fig. 4m), they also appear to be more impacted by alping than the first lactation cows as highlighted by negative Δd-values (Fig. 4o). The influence of pregnancy stage appears to affect milk production during alping, especially for cows calving in autumn (Fig 4p and 4r). Further, higher cows and/or with steeper foot angle produce more milk both before and during alping than lower ones with gentle foot angle (Fig. 4s, t, v, w). However, alping appears to negatively impact such cows, especially higher ones (Fig. 4u and 4x).

## Discussion

### The importance of calving season

The proposed model succeeded in quantifying the impact of alping on milk production by assuming a Wilmink pattern for cows experiencing the same conditions (Fig.2; [40]). As expected, total milk production resulted globally higher for cows with alping occurring at the end of the lactation, since the drop in production happens later in the cycle. Anyway, winter calving might still be financially attractive for farmers since milk produced in the alps will have a higher economic value on the market and productivity will be higher during alping (Fig. 3).

Calving season also influences the way an animal is prompt to respond to environmental stress, with a greater impact of transhumance (i.e. greater d-parameter in absolute value) for cows calving in autumn and therefore alping at the end of their lactation cycle. Increased feed intake is known to have distinct effects on milk production depending on the lactation stage [44], and from what we observe it appears that milk production at the end of the lactation cycle is more sensitive to environmental changes. Similarly, when studying the effect of the considered factors, we showed that the between-group difference in milk production during alping is almost always greater for autumn calving.

### Effect of the environment and climate change

Climate change requires species to adapt quickly to new and extreme climatic conditions [45]. In this context, cattle survival and annexed services for humans are threatened because of the low adaptive potential observed for industrial breeds [46]. In Switzerland, climatic conditions are becoming hotter and dryer [47], which exhorts to better understand the effects of climate on cattle welfare and production both at farms and during transhumance. Here, we observe a sensible negative effect of precipitation in spring (Fig 4e-f), probably because of their influence on forage growth [31]. Interestingly, heat waves (which are known to highly affect cattle productivity [48]) were found to have minimal impact on milk production during alping, probably because temperatures at high altitude rarely reach problematic thresholds. Similarly, cold spells seem to have an almost negligible influence (Sup. Mat. S1). The observed effect of biogeographical regions on production can be explained by the difference in spring precipitation between such regions (158mm/month versus 98mm/month for the Northern flank and the Eastern part, respectively). Altitude confirmed its effect on productivity [23], being intrinsically connected with climatic conditions and vegetation type.

### Effect of physiological and morphological factors

Lactation number has long been known to strongly influence milk production [49], and this also holds for milk production during alping (Fig 4m-o). Even more important, pregnancy stage was found to have a significant impact on milk production during alping, especially when calving occurs in autumn (Fig. 4p-r). In order to optimize milk yield, cows are generally inseminated a few months after calving, to reach a time span of one year between lactation cycles, implying pregnancy stage not to be considered in lactation models to avoid strong collinearity with calving season [15,16]. However, correlation among these variables was not extreme in the present case (r^2^=0.8), most likely because of unsuccessful inseminations leading some cows to delay pregnancy. These results must be interpreted with care, as cows with an early pregnancy are prone to fertility problems

Many recent research effort focused on increasing yield in cattle, leading to augmented cattle size [50] but disregarding important side-effects such as the loss of adaptive traits through genetic erosion [51]. This phenomenon might become deleterious for transhumance. For instance, despite showing higher productive performances even at alping, higher cow appear to be more impacted when moved to high mountain pastures (Fig. 4s-u). As for foot angler, steep angle is associated with a smaller risk of developing hoof diseases [52]. Cows with steeper foot angle were observed to produce more milk both in lowland farm and during alping, but this factor appears to be have limited on the d-parameter (Fig. 4v-x).

### Limitations

Traditionally, lactation modelling is performed on an individual basis, and usually relies on daily or weekly milk records [53]. Here, we based our work on a database composed of monthly milk records, which required the transformation of the data into daily averages over thousands of cows to avoid over-parameterisation in the model. This averaging might have diluted the strength of the effect we investigated.

Moreover, the proposed approach still misses validation, which could be achieved by relying on individual observations recorded daily or weekly and belonging to different breeds from the one used here.

Next, the amount of observations among calving months was not constant in the dataset, which possibly made the estimates from the winter months less robust. Further, a hidden age effect – as older cows tend to calf later in the season-could have biased the observed differences in milk productions among groups.

Last but not least, the model does not explicitly take into account cow feeding during alping, which is likely to affect milk production [24]. Indeed, the use of concentrate feeding varies among alps and among cows of the same alp. Particularly, differences in milk yield with different calving season could be globally influenced by varying concentrates feeding, with cows at an early stage in the lactation cycle – and thus producing a substantial amount of milk - potentially receiving more concentrates.

## 6. Conclusion

Transhumance is a traditional farming practice which supports the preservation of both agricultural biodiversity and the socio-cultural heritage of human communities. Nevertheless, a loss in productivity is typically linked with alped livestock, which might discourage farmers from pursuing transhumance and poses its beneficial side-effects on ecosystems under threat. Here, we combined biological, geo-environmental and computer science tools to better understand the influence of environmental, physiological and morphological factors on milk productivity during transhumance. We relied on high resolution meteorological data and five millions georeferenced monthly milk records as collected from over 200,000 Braunvieh cows in Switzerland. We show that both environmental and morphological factors have limited influence on animal production, with dry conditions in spring being nevertheless the most affecting environmental factor. This evidence suggests that animal production during transhumance might become even more insecure in future years due to climate change, and stress therefore the urgency of devising strategies to protect this practice. On the other hand, physiological factors such as lactation number, pregnancy stage have strong impact on milk production during the whole lactation cycle.

## Supporting information

Sup. Mat. 1

## Acknowledgments

We are grateful to the breeding organisation Braunvieh Schweiz for extracting and distributing the full data set from their database

## Ethical Statement

No ethical statement to declare.

## Funding Statement

No funding source to declare.

## Data Accessibility

The data was provided from the Braunvieh-CH association, under the explicit conditions that it will not be shared nor used for other studies. However, a partial dataset is available with the average milk production during alping from 20000 cows, together with lactation information (calving date, lactation number) and environmental data at the location of alping. Cows were chosen randomly, with equal number of animals per year, lactation number and calving month. Furthermore, researchers interested in performing studies on these data may contact directly the association (see contact information homepage.braunvieh.ch). The code used for this article is available at https://github.com/SolangeD/lactModel

## Competing Interests

We have no competing interests.

## Authors’ Contributions

SD performed most of the analyses with the help of AB. CF and SJ supervised the work. SD wrote the first draft of the article and all authors contributed in improving this draft.

## Notes

https://datadryad.org/stash/share/ClYGHz9eJxS1DmE5UWl4reOM0sMVQhH_F3Prt0Htttg

### Competing Interest Statement

The authors have declared no competing interest.

## References

1. Bunce RGH, Pérez-Soba M, Smith M. 2009 Assessment of the extent of agroforestry systems in Europe and their role within transhumance systems. In Agroforestry in Europe, pp. 321–329. Springer.

2. Olea PP, Mateo-Tomás P. 2009 The role of traditional farming practices in ecosystem conservation: the case of transhumance and vultures. Biological conservation 142, 1844–1853.

3. Liechti K, Biber J-P. 2016 Pastoralism in Europe: characteristics and challenges of highland-lowland transhumance: -EN- -FR-Le pastoralisme en Europe: caractéristiques et défis de la transhumance de la montagne vers la plaine-ES-El pastoreo en Europa: características y problemas de la trashumancia de tierras altas-tierras bajas. Rev. Sci. Tech. OIE 35, 561–575. (doi:10.20506/rst.35.2.2541)

4. Herzog F, Bunce RG, Pérez-Soba M, Jongman RH, Sal AG, Austad I. 2005 Policy options to support transhumance and biodiversity in European mountains: a report on the TRANSHUMOUNT Stakeholder Workshop, Landquart/Zurich, Switzerland, 26-28 May 2004. Mountain Research and Development 25, 82–84.

5. Gellrich M, Zimmermann NE. 2007 Investigating the regional-scale pattern of agricultural land abandonment in the Swiss mountains: a spatial statistical modelling approach. Landscape and Urban Planning 79, 65–76.

6. Gellrich M, Baur P, Koch B, Zimmermann NE. 2007 Agricultural land abandonment and natural forest re-growth in the Swiss mountains: A spatially explicit economic analysis. Agriculture, Ecosystems & Environment 118, 93–108. (doi:10.1016/j.agee.2006.05.001)

7. Newesely C, Tasser E, Spadinger P, Cernusca A. 2000 Effects of land-use changes on snow gliding processes in alpine ecosystems. Basic and Applied Ecology 1, 61–67.

8. Lauber S. 2013 Avenir de l’économie alpestre suisse. Faits, analyses et éléments de réflexion issus du programme de recherche AlpFUTUR. Institut fédéral de recherche sur la forêt, la neige et le paysage (WSL), Birmensdorf

9. In press. Agrarbericht 2019-Sömmerungsbetriebe. See https://www.agrarbericht.ch/de/betrieb/strukturen/soemmerungsbetriebe (accessed on 13 February 2020).

10. Jeretina J, Babnik D, Skorjanc D. 2013 Modeling lactation curve standards for test-day milk yield in Holstein, Brown Swiss and Simmental cows. The J Anim and Plant Sci 233, 754–62.

11. Wood PDP. 1967 Algebraic Model of the Lactation Curve in Cattle. Nature 216, 164. (doi:10.1038/216164a0)

12. Zendri F, Ramanzin M, Bittante G, Sturaro E. 2016 Transhumance of dairy cows to highland summer pastures interacts with breed to influence body condition, milk yield and quality. Italian Journal of Animal Science 15, 481–491.

13. Cassandro M, Comin A, Ojala M, Zotto RD, De Marchi M, Gallo L, Carnier P, Bittante G. 2008 Genetic Parameters of Milk Coagulation Properties and Their Relationships with Milk Yield and Quality Traits in Italian Holstein Cows. Journal of Dairy Science 91, 371–376. (doi: 10.3168/jds.2007-0308)

14. Jõudu I, Henno M, Kaart T, Püssa T, Kärt O. 2008 The effect of milk protein contents on the rennet coagulation properties of milk from individual dairy cows. International Dairy Journal 18, 964–967. (doi:10.1016/j.idairyj.2008.02.002)

15. Tekerli M, Akinci Z, Dogan I, Akcan A. 2000 Factors affecting the shape of lactation curves of Holstein cows from the Balikesir province of Turkey. Journal of Dairy Science 83, 1381–1386.

16. Wilmink JBM. 1987 Adjustment of test-day milk, fat and protein yield for age, season and stage of lactation. Livestock Production Science 16, 335–348.

17. Hayes BJ, Carrick M, Bowman P, Goddard ME. 2003 Genotype$\times$ environment interaction for milk production of daughters of Australian dairy sires from test-day records. Journal of dairy science 86, 3736–3744.

18. Hahn GL. 1999 Dynamic responses of cattle to thermal heat loads. Journal of Animal Science 77, 10.

19. Bryant JR, López-Villalobos N, Pryce JE, Holmes CW, Johnson DL. 2007 Quantifying the effect of thermal environment on production traits in three breeds of dairy cattle in New Zealand. New Zealand Journal of Agricultural Research 50, 327–338.

20. Gilgen AK, Buchmann N. 2009 Response of temperate grasslands at different altitudes to simulated summer drought differed but scaled with annual precipitation. Biogeosciences 6, 2525–2539.

21. Signarbieux C, Feller U. 2008 Effects of an extended drought period on grasslands at various altitudes in Switzerland: A field study. In Photosynthesis. Energy from the Sun, pp. 1371–1374. Springer.

22. Jurt C, Häberli I, Rossier R. 2015 Transhumance Farming in Swiss Mountains: Adaptation to a Changing Environment. Mountain Research and Development 35, 57–65.

23. Gorlier A, Lonati M, Renna M, Lussiana C, Lombardi G, Battaglini LM. 2013 Changes in pasture and cow milk compositions during a summer transhumance in the western Italian Alps. Journal of Applied Botany and Food Quality 85, 216.

24. Leiber F, Kreuzer M, Leuenberger H, Wettstein HR. 2006 Contribution of diet type and pasture conditions to the influence of high altitude grazing on intake, performance and composition and renneting properties of the milk of cows. Animal Research 55, 37–53.

25. Bovolenta S, Ventura W, Malossini F. 2002 Dairy cows grazing an alpine pasture: effect of pattern of supplement allocation on herbage intake, body condition, milk yield and coagulation properties. Animal Research 51, 15–23.

26. Berry NR, Sutter F, Bruckmaier RM, Blum JW, Kreuzer M. 2001 Limitations of high Alpine grazing conditions for early-lactation cows: effects of energy and protein supplementation. Animal Science 73, 149–162.

27. Horn M, Steinwidder A, Starz W, Pfister R, Zollitsch W. 2014 Interactions between calving season and cattle breed in a seasonal Alpine organic and low-input dairy system. Livestock Science 160, 141–150. (doi:10.1016/j.livsci.2013.11.014)

28. Horn M, Steinwidder A, Gasteiner J, Podstatzky L, Haiger A, Zollitsch W. 2013 Suitability of different dairy cow types for an Alpine organic and low-input milk production system. Livestock Science 153, 135–146.

29. Bertaglia M, Joost S, Roosen J. 2007 Identifying European marginal areas in the context of local sheep and goat breeds conservation: A geographic information system approach. Agricultural Systems 94, 657–670. (doi:10.1016/j.agsy.2007.02.006)

30. ICAR. 2014 ICAR Recording Guidelines approved by the General Assembly held in Berlin, Germany, on May 2014.

31. Jonas T, Rixen C, Sturm M, Stoeckli V. 2008 How alpine plant growth is linked to snow cover and climate variability. Journal of Geophysical Research: Biogeosciences 113.

32. Hayes B. 2013 Overview of statistical methods for genome-wide association studies (GWAS). Genome Wide Association Studies and Genomic Prediction, 149–169.

33. Olori VE, Brotherstone S, Hill WG, McGuirk BJ. 1997 Effect of gestation stage on milk yield and composition in Holstein Friesian dairy cattle. Livestock Production Science 52, 167–176.

34. Ugurlu M, Teke B, Akdag F, Arslan S. 2014 EFFECT OF TEMPERATUREHUMIDITY INDEX, COLD STRESS INDEX AND DRY PERIOD LENGHT ON BIRTH WEIGHT OF JERSEY CALF. Bulgarian Journal of Agricultural Science 20, 1227–1232.

35. Luo W, Taylor MC, Parker SR. 2008 A comparison of spatial interpolation methods to estimate continuous wind speed surfaces using irregularly distributed data from England and Wales. International journal of climatology 28, 947–959.

36. Swisstopo. In press. DHM25/200.

37. FOEN. 2001 Die biogeographischen Regionen der Schweiz.

38. Meteoswiss. 2018 The climate of Switzerland.

39. Val-Arreola D, Kebreab E, Dijkstra J, France J. 2004 Study of the Lactation Curve in Dairy Cattle on Farms in Central Mexico. Journal of Dairy Science 87, 3789–3799. (doi:10.3168/jds.S0022-0302(04)73518-3)

40. Macciotta NPP, Vicario D, Cappio-Borlino A. 2005 Detection of different shapes of lactation curve for milk yield in dairy cattle by empirical mathematical models. Journal of Dairy Science 88, 1178–1191.

41. R Core Team. 2018 R: A Language and Environment for Statistical Computing. Vienna, Austria: R Foundation for Statistical Computing. See https://www.R-project.org/.

42. Bonferroni CE, Bonferroni C, Bonferroni CE. 1936 Teoria statistica delle classi e calcolo delle probabilita’.

43. Zeileis A, Hothorn T. 2002 Diagnostic checking in regression relationships.

44. Johnson CL. 1984 The effect of feeding in early lactation on feed intake, yields of milk, fat and protein and on live-weight change over one lactation cycle in dairy cows. The Journal of Agricultural Science 103, 629–637.

45. Hayes BJ, Bowman PJ, Chamberlain AJ, Savin K, Van Tassell CP, Sonstegard TS, Goddard ME. 2009 A validated genome wide association study to breed cattle adapted to an environment altered by climate change. PLoS One 4, e6676.

46. Taberlet P, Valentini A, Rezaei HR, Naderi S, Pompanon F, Negrini R, Ajmone-Marsan P. 2008 Are cattle, sheep, and goats endangered species? Mol. Ecol. 17, 275–284.(doi:10.1111/j.1365-294X.2007.03475.x)

47. IPCC. 2014 Climate Change 2014 Synthesis report; A report of the intergovernmental panel on climate change.

48. Kadzere CT, Murphy MR, Silanikove N, Maltz E. 2002 Heat stress in lactating dairy cows: a review. Livestock production science 77, 59–91.

49. Strucken EM, Bortfeldt RH, De Koning DJ, Brockmann GA. 2012 Genome-wide associations for investigating time-dependent genetic effects for milk production traits in dairy cattle. Animal genetics 43, 375–382.

50. Tsuruta S, Misztal I, Lawlor TJ. 2004 Genetic Correlations Among Production, Body Size, Udder, and Productive Life Traits Over Time in Holsteins. Journal of Dairy Science 87, 1457–1468. (doi:10.3168/jds.S0022-0302(04)73297-X)

51. Notter DR. 1999 The importance of genetic diversity in livestock populations of the future. J ANIM SCI 77, 61–69.

52. Rogers GW. 2002 ASPECTS OF MILK COMPOSITION, PRODUCTIVE LIFE AND TYPE TRAITS IN RELATION TO MASTITIS AND OTHER DISEASES IN DAIRY CATTLE., 7.

53. Olori VE, Brotherstone S, Hill WG, McGuirk BJ. 1999 Fit of standard models of the lactation curve to weekly records of milk production of cows in a single herd. Livestock Production Science 58, 55–63.

