## Supplementary material for "Big dairy data to disentangle the effect of geo-environmental, physiological and morphological factors on milk production of mountain-pastured Braunvieh cows": Sup. Mat. 1

| **Crit** | **Calving month** | **Δmilk alp** | **Δmilk total** | **Δd** | **p-value** |
| --- | --- | --- | --- | --- | --- |
| Lact # | 9 | 5.3 | 16.9 | 0.0070 | 1.16E-01 |
|  | 10 | 7.0 | 17.5 | -0.0036 | 5.74E-01 |
|  | 11 | 11.3 | 18.8 | -0.0071 | 4.55E-04 |
|  | 12 | 16.2 | 19.8 | -0.0077 | 1.69E-03 |
|  | 1 | 18.9 | 19.4 | -0.0066 | 4.61E-01 |
|  | 2 | 21.4 | 19.8 | -0.0080 | 1.00E+00 |
| Pregnancy stage | 9 | 23.2 | 2.5 | 0.0333 | 2.16E-15 |
|  | 10 | 18.4 | 4.3 | 0.0278 | 1.98E-20 |
|  | 11 | 12.1 | 4.1 | 0.0250 | 1.40E-27 |
|  | 12 | 6.4 | 3.6 | 0.0203 | 6.84E-18 |
|  | 1 | 2.5 | 2.9 | 0.0161 | 1.63E-04 |
|  | 2 | 1.0 | 2.8 | 0.0202 | 1.93E-01 |
| THI-3d | 9 | 3.2 | 0.2 | 0.0065 | 1.78E-01 |
|  | 10 | 1.8 | 0.3 | 0.0039 | 1.17E-01 |
|  | 11 | 1.3 | 0.4 | 0.0030 | 1.28E-01 |
|  | 12 | 0.1 | 0.1 | 0.0004 | 1.00E+00 |
|  | 1 | 0.8 | 0.6 | 0.0023 | 5.38E-01 |
|  | 2 | 1.5 | 1.7 | 0.0046 | 4.96E-02 |
| THI-30d | 9 | 0.1 | 0.0 | 0.0002 | 1.00E+00 |
|  | 10 | 2.3 | 0.3 | 0.0049 | 3.57E-02 |
|  | 11 | 1.9 | 0.5 | 0.0044 | 5.24E-03 |
|  | 12 | 0.8 | 0.4 | 0.0022 | 3.10E-01 |
|  | 1 | 0.6 | 0.5 | 0.0018 | 1.00E+00 |
|  | 2 | 1.0 | 1.1 | 0.0030 | 5.30E-01 |
| CSI-3d | 9 | -1.3 | -0.1 | -0.0026 | 1.00E+00 |
|  | 10 | -0.1 | 0.0 | -0.0002 | 1.00E+00 |
|  | 11 | -0.7 | -0.2 | -0.0017 | 1.00E+00 |
|  | 12 | -0.4 | -0.2 | -0.0011 | 1.00E+00 |
|  | 1 | -0.5 | -0.4 | -0.0015 | 1.00E+00 |
|  | 2 | -0.8 | -0.9 | -0.0023 | 9.23E-01 |
| CSI-30d | 9 | -0.7 | 0.0 | -0.0014 | 1.00E+00 |
|  | 10 | -0.2 | 0.0 | -0.0004 | 1.00E+00 |
|  | 11 | 0.2 | 0.1 | 0.0005 | 1.00E+00 |
|  | 12 | 0.2 | 0.1 | 0.0004 | 1.00E+00 |
|  | 1 | -0.6 | -0.5 | -0.0018 | 9.44E-01 |
|  | 2 | -1.4 | -1.6 | -0.0043 | 7.45E-02 |
| Precipitations in sping | 9 | 9.3 | 0.5 | 0.0189 | 3.77E-16 |
|  | 10 | 8.4 | 1.2 | 0.0181 | 1.22E-32 |
|  | 11 | 7.2 | 2.0 | 0.0171 | 6.47E-44 |
|  | 12 | 6.0 | 2.9 | 0.0159 | 3.93E-39 |
|  | 1 | 5.5 | 4.2 | 0.0159 | 2.09E-25 |
|  | 2 | 5.1 | 5.8 | 0.0157 | 1.52E-19 |

| **Crit** | **Calving month** | **Δmilk alp** | **Δmilk total** | **Δd** | **p-value** |
| --- | --- | --- | --- | --- | --- |
| Biogeographical region | 9 | 10.4 | 0.6 | 0.0209 | 3.82E-20 |
|  | 10 | 9.2 | 1.3 | 0.0198 | 9.38E-47 |
|  | 11 | 8.2 | 2.3 | 0.0196 | 1.69E-73 |
|  | 12 | 6.9 | 3.3 | 0.0183 | 3.84E-59 |
|  | 1 | 5.3 | 4.0 | 0.0152 | 4.47E-19 |
|  | 2 | 5.1 | 5.9 | 0.0159 | 2.37E-11 |
| Altitude | 9 | 5.3 | 0.3 | 0.0107 | 6.72E-07 |
|  | 10 | 5.3 | 0.7 | 0.0115 | 1.18E-18 |
|  | 11 | 5.6 | 1.6 | 0.0133 | 1.35E-44 |
|  | 12 | 4.6 | 2.2 | 0.0121 | 1.10E-37 |
|  | 1 | 3.6 | 2.8 | 0.0104 | 2.57E-14 |
|  | 2 | 3.7 | 4.2 | 0.0115 | 5.20E-12 |
| Difference in altitude | 9 | 1.2 | 0.1 | 0.0025 | 1.00E+00 |
|  | 10 | 1.1 | 0.2 | 0.0023 | 3.30E-01 |
|  | 11 | 2.2 | 0.6 | 0.0052 | 2.27E-07 |
|  | 12 | 2.6 | 1.3 | 0.0070 | 2.14E-13 |
|  | 1 | 3.5 | 2.7 | 0.0101 | 3.84E-15 |
|  | 2 | 3.8 | 4.3 | 0.0118 | 2.59E-13 |
| Aspect (100m) | 9 | 0.4 | 0.0 | 0.0008 | 1.00E+00 |
|  | 10 | 0.8 | 0.1 | 0.0018 | 6.76E-01 |
|  | 11 | 0.3 | 0.1 | 0.0006 | 1.00E+00 |
|  | 12 | 0.3 | 0.1 | 0.0007 | 1.00E+00 |
|  | 1 | 0.3 | 0.2 | 0.0008 | 1.00E+00 |
|  | 2 | -0.2 | -0.3 | -0.0008 | 1.00E+00 |
| Aspect (1km) | 9 | 1.6 | 0.1 | 0.0032 | 5.04E-01 |
|  | 10 | 1.1 | 0.2 | 0.0025 | 1.78E-01 |
|  | 11 | 0.8 | 0.2 | 0.0020 | 7.99E-02 |
|  | 12 | 0.7 | 0.4 | 0.0020 | 8.61E-02 |
|  | 1 | 0.5 | 0.4 | 0.0015 | 1.00E+00 |
|  | 2 | 0.4 | 0.4 | 0.0012 | 1.00E+00 |
| Height | 9 | 7.7 | 14.9 | -0.0137 | 7.31E-04 |
|  | 10 | 8.5 | 14.5 | -0.0125 | 4.68E-06 |
|  | 11 | 9.9 | 14.7 | -0.0091 | 3.89E-04 |
|  | 12 | 11.5 | 14.7 | -0.0089 | 9.68E-03 |
|  | 1 | 11.2 | 12.4 | -0.0022 | 1.00E+00 |
|  | 2 | 11.9 | 10.9 | 0.0105 | 9.12E-01 |
| Foot angle | 9 | 2.8 | 6.6 | -0.0095 | 1.32E-02 |
|  | 10 | 3.3 | 6.2 | -0.0079 | 2.60E-03 |
|  | 11 | 4.8 | 7.2 | -0.0082 | 3.21E-03 |
|  | 12 | 7.4 | 8.4 | -0.0065 | 4.39E-01 |
|  | 1 | 9.6 | 9.3 | -0.0025 | 1.00E+00 |
|  | 2 | 11.1 | 9.4 | -0.0005 | 1.00E+00 |

Sup. Mat. For each criterion and calving month, the between-group difference in milk production in the alp and over the total lactation cycle is reported, together with the Δd (reflecting how differently the two groups are impacted by alping) and its associated significance.
